# A new family of neural wiring receptors across bilaterians defined by phylogenetic, biochemical and structural evidence

**DOI:** 10.1101/462036

**Authors:** Shouqiang Cheng, Yeonwoo Park, Justyna D. Kurleto, Mili Jeon, Kai Zinn, Joseph W. Thornton, Engin Özkan

## Abstract

The evolution of complex nervous systems was accompanied by the expansion of groups of protein families, most notably cell adhesion molecules, surface receptors and their ligands. These proteins mediate axonal guidance, synapse targeting, and other neuronal wiring-related functions. Recently, members of a set of thirty interacting cell surface proteins belonging to two newly defined families of the immunoglobulin superfamily (IgSF) in fruit flies were discovered to label different subsets of neurons in the brain and ventral nerve cord. They have been shown to be involved in synaptic targeting and morphogenesis, retrograde signaling, and neuronal survival. Here we show that these proteins, denoted as Dprs and DIPs, belong to a family of two and three-Ig domain molecules in bilaterians generally known for neuronal wiring functions. In protostomes, the ancestral Dpr/DIP gene has duplicated to form heterophilic partners, such as Dprs and DIPs, while in deuterostomes, they have evolved to create the IgLON family of neuronal receptors. In support of this phylogeny, we show that IgLONs interact with each other, and that their complexes can be broken by mutations designed using homology models based on Dpr and DIP structures. Similarly, the nematode orthologs ZIG-8 and RIG-5 can form heterophilic and homophilic complexes structurally matching Dpr-DIP and DIP-DIP complexes. The evolutionary, biochemical and structural relationships we demonstrate here provides insights into neural development and the rise of complexity in metazoans.

**Significance Statement:** Cell surface receptors assign and display unique identities to neurons, and direct proper and robust wiring of neurons to create functional neural circuits. Recent work has identified two new classes of receptors in fruit flies, called the Dpr and DIP families with 30 members, which interact in 38 pairwise combinations. These proteins are implicated in neural identity, wiring and survival in many parts of the fly nervous system. Here, using evolutionary, biochemical and structural evidence, we show that Dprs and DIPs are members of an ancient bilaterian family of receptors. Members of this family share functional roles relevant to wiring across species, and are likely crucial in the emergence of the bilaterian nervous systems common to vertebrate and invertebrate animals.

## INTRODUCTION

Immunoglobulin superfamily (IgSF) proteins, which form the largest single-pass cell surface and adhesion family in humans, are crucial to animal development, and have gone through large expansions as complexity in metazoans increased (1–3). They have been studied in both vertebrate and invertebrate systems in the context of development and function of the immune and nervous systems. Unlike in the immune system, neural functions, such as neurite outgrowth, guidance, fasciculation, and synaptic targeting, employ IgSF and other cell surface molecules that are usually conserved between vertebrates and invertebrates. As the central functionality of IgSF proteins on the cell surface is mediated through the recognition of IgSF or other surface receptors and ligands, recent efforts have focused on deorphanization of these proteins in vertebrates (4, 5) and invertebrates (6) through high-throughput interactome studies. However, genomic and interactomic data can be difficult to interpret, especially when proteins are not annotated for function and orthologous protein present in vertebrate and invertebrate model organisms cannot be identified.

Our interactome studies on *Drosophila* IgSF proteins have recently revealed two protein families with remarkable neural expression patterns: the Dpr family, named after the founding member Defective proboscis extension response (7), and their binding partners, the Dpr-interacting proteins (DIPs) (6, 8). Dprs and DIPs form a complex interaction network consisting of 38 interactions among 30 proteins. Most of the Dprs and DIPs that have been studied thus far are expressed exclusively in the nervous system. In the pupal optic lobe, the larval ventral nerve cord, and the neuromuscular system, each Dpr and DIP is expressed in a unique subset of neurons (6, 8, 9). One Dpr is also expressed in postsynaptic muscle cells (11). In the optic lobe, Dprs and DIPs are expressed in distinctive combinations in different neuronal types, and synaptic targeting defects and neuronal death have been observed in *dpr11* and *DIP-γ* mutants (8, 9). In the neuromuscular system, *dpr11* and *DIP-γ* mutants show synapse maturation defects, while Dpr10 and DIP-α are shown to be necessary for the formation of synapses onto specific muscle targets (10, 11). In the olfactory system, Dprs and DIPs express in unique combinatorial patterns in olfactory receptor neurons and are necessary for neuronal adhesion and glomerulus formation (12). Overall, the available data suggest that Dprs and DIPs serve neuronal wiring functions, likely by acting as “identification tags” for neurons, and physically guiding their connectivity. Since the numbers of known synaptic targeting molecules are limited, the study of Dprs and DIPs has the potential to illuminate mechanisms involved in development of synaptic circuits across the nervous system in *Drosophila*.

Dprs and DIPs have domain structures that are similar to those of many other IgSF proteins (3). Following a signal peptide, they carry two and three immunoglobulin (Ig) domains, respectively. Dprs and DIPs interact via their N-terminal Ig domains, creating a pseudosymmetric Ig-Ig complex (8). The C-terminal ends of Dprs and DIPs are strongly hydrophobic, either serving as transmembrane helices or as recognition sites for glycosylphosphatidylinositol (GPI) linkages to the plasma membrane. Most Dprs and DIPs do not have intracellular domains, and lack conserved features in their juxtamembrane regions.

While Dpr and DIP orthologs can be identified in arthropod species, it is not clear whether they exist in other animals. Furthermore, it is not known why Dprs and DIPs have gone through a large expansion in arthropods, and if their orthologs have gone through similar processes. Finally, whether the non-arthropod orthologs, if they exist, function in neural development with similar or overlapping roles has not been explored. Therefore, we set out to uncover how Dprs and DIPs expanded in the arthropod line, reveal their orthologs across major metazoan groupings, and establish biochemical and structural similarities among the proposed proteins. Here, we show that Dprs and DIPs have originated from a common ancestor with vertebrate proteins known as IgLONs, a family of five neuronal proteins in humans. The shared roles of these proteins in neurite outgrowth and synapse formation suggest that the family consisting of these proteins, here named *Wirins*, is primarily involved in nervous system development across bilaterians. The Wirin family expanded independently in vertebrates, arthropods, and mollusks through multiple gene duplications. We further show that the homophilic and heterophilic interactions characteristic of Dprs and DIPs are also observed among vertebrate IgLONs and the nematode orthologs. In addition, we demonstrate that the molecular interfaces known to mediate Dpr-DIP interactions are used by the orthologs as well. Overall, we describe a family of proteins that share conserved roles in nervous system development, which have evolved early in bilaterians, and have undergone gene expansions in conjunction with the evolution of complex nervous systems.

## RESULTS

### The Wirin family of IgSF proteins with neural wiring functions

Dprs and DIPs were discovered and characterized in *Drosophila melanogaster*. To identify their homologs in other organisms, we carried out a phylogenetic analysis. For sequence mining, a putative Dpr or DIP homolog was defined as a protein that shows the highest sequence similarity to a Dpr or DIP among all proteins in *D. melanogaster*. In this way, distant IgSF homologs could be excluded from analysis. All examined protostomes had both Dpr and DIP homologs. Chordates had a family of proteins called IgLONs that appeared as DIP homologs. Non-chordate deuterostomes and non-bilaterians lacked recognizable Dpr or DIP homologs. Using human IgLONs as reference, additional IgLON homologs were identified in protostomes: CG34353, CG7166, Klingon, Lachesin, and Wrapper. No additional homologs for these proteins could be found in chordates.

Figure 1 shows the maximum likelihood (ML) phylogeny of the above proteins. Four IgSF families – Nectin, Necl, Kirrel, and Nephrin – were included to help root the phylogeny (See *supplemental information* for family definitions). There were multiple equally parsimonious ways to place the root, all implying the same number of gene gains and losses. However, in all cases, CG34353, CG7166, DIPs, Dprs, IgLONs, Klingon, and Lachesin formed a monophyletic group. We refer to this group as the Wirin (*Wir*ing *I*mmunoglobuli*n*) family. The Wirin family began as a single gene in the last common ancestor of bilaterians. It evolved into the IgLON subfamily in deuterostomes. In protostomes, it evolved into the DIP, Dpr, Klingon, and Lachesin subfamilies through serial gene duplications. IgLONs are thus co-orthologous to DIPs, Dprs, Klingons, and Lachesins. This phylogeny, along with the known functions of DIPs, Dprs, and IgLONs, indicates a conserved role for the Wirin family as neuronal wiring proteins in the bilaterian nervous system.

**Fig. 1.**
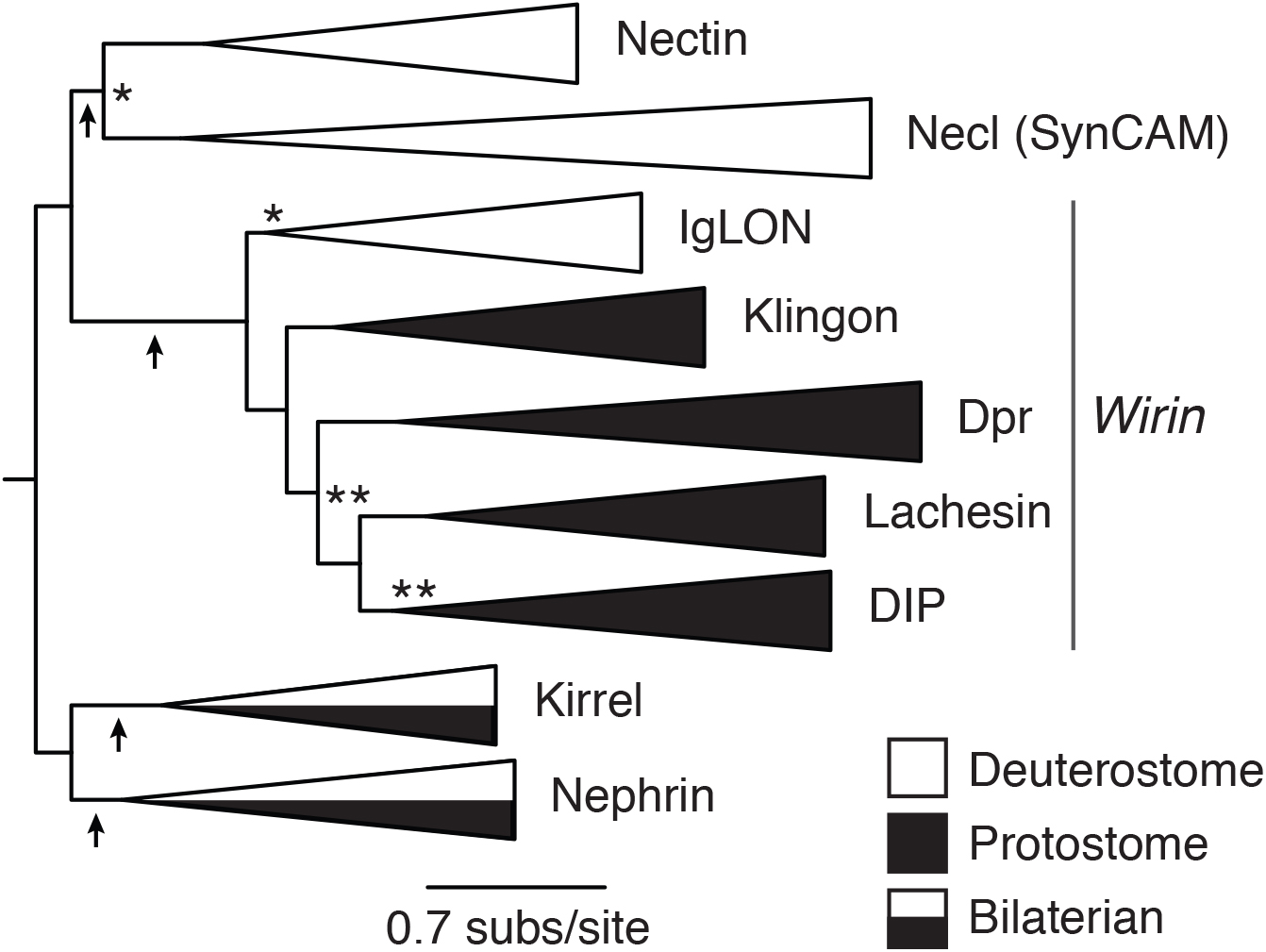
The maximum likelihood (ML) phylogeny of the Wirin family. Approximate likelihood ratio statistics (aLRS) are shown as branch supports: ** < 9.2 (= 2 ln100), * < 3.2 (= 2 ln5). Unmarked branches have aLRS > 9.2. The arrows indicate alternative ways of rooting the phylogeny that imply the same number of gene gains and losses as the current phylogeny.

Each Wirin subfamily expanded through subsequent gene duplications. Figure 2A shows the ML phylogeny of the IgLON subfamily. When clades subtending to poorly supported branches are rearranged (the more basal cyclostome clade and the chondrichthyan NTM/OBCAM), the phylogeny becomes congruent with gene family expansion through the two genome duplications in chordate evolutionary history. An additional gene duplication in tetrapods gave rise to total five IgLONs in human. The Dpr and DIP subfamilies expanded through a dozen or more gene duplications independently in arthropods and mollusks (Fig. S1 and S2). Other protostomes, such as annelids, brachiopods, nematodes, and tardigrades, remained with one or two paralogs. The Klingon subfamily expanded in hexapods, resulting in four paralogs in *D. melanogaster* (CG34353, CG7166, Klingon, and Wrapper; Fig. S3A). The Lachesin subfamily expanded in lophotrochozoans, giving rise to two paralogs in mollusks and four paralogs in annelids (Fig. S3B).

**Fig. 2.**
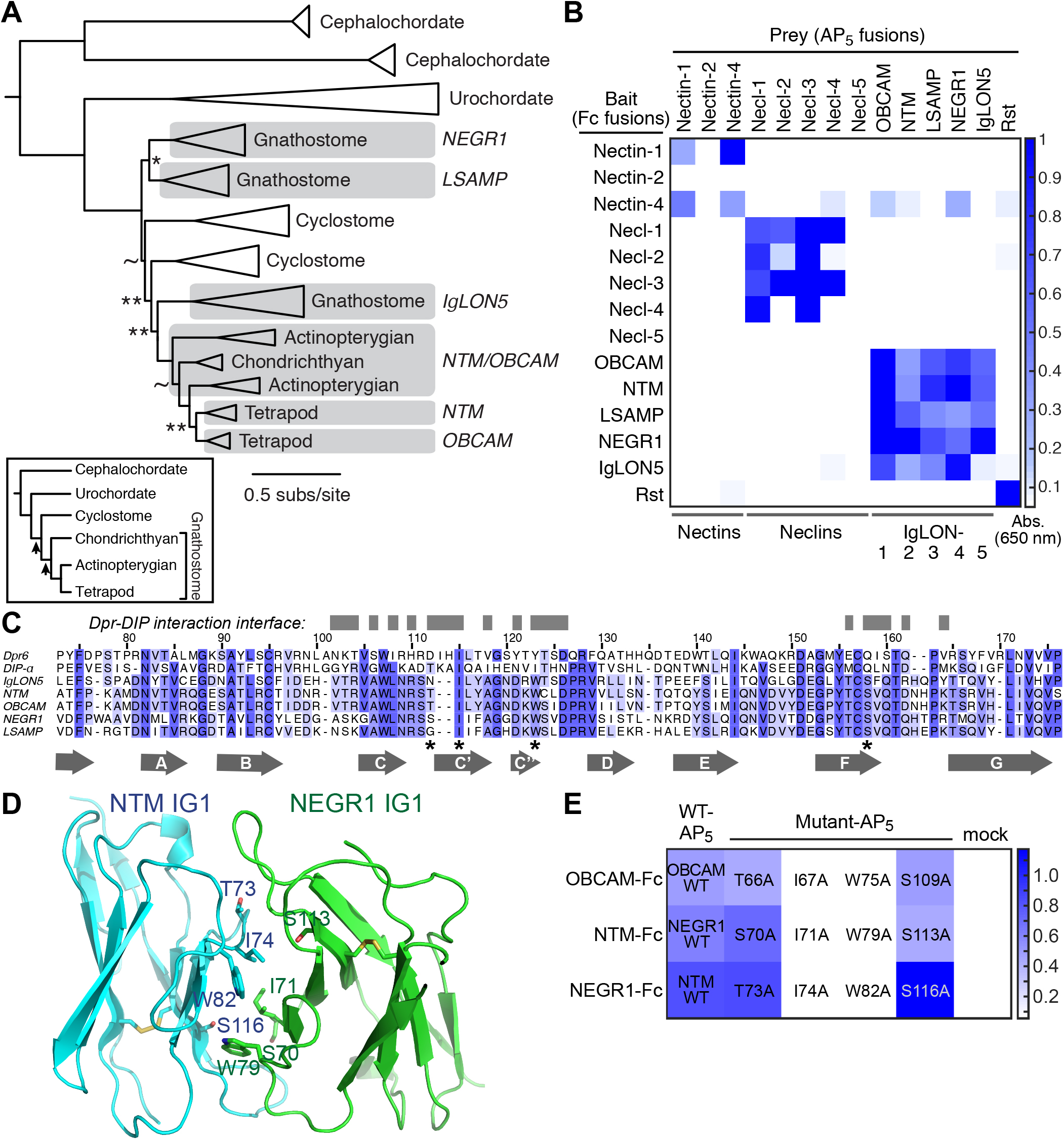
IgLONs are the vertebrate Wirins. **A.** The ML phylogeny of the IgLON family. Approximate likelihood ratio statistics (aLRS) are shown as branch supports: ** < 9.2 (= 2 ln100), * < 4.6 (= 2 ln10), ~ < 2.2 (= 2 ln3). Unmarked branches have aLRS > 9.2. The inset shows the chordate phylogeny with arrows marking the two genome duplications. **B.** Binding experiments for mouse IgLONs, Nectins and Nectin-like proteins using ECIA. The homodimeric Rst, the fly ortholog of mammalian Kirrels, serves as a positive control. **C.** Sequence alignment of the IG1 domains of mouse IgLONs, Dpr6 and DIP-α. Amino acids known to be at the Dpr-DIP interface, defined by a 4 Å cutoff of the binding partner, are labeled as gray blocks above the alignment. Amino acids mutated in Fig. 2E are labeled with an asterisk. Sequence numbers above are for Dpr6. **E.** Mutations at the predicted interfaces of the OBCAM-OBCAM and NTM-NEGR1 complexes affect binding as tested using ECIA. To effectively compare wild-type (WT) to mutants, protein concentrations within each mutant series were normalized.

### IgLONs interact with each other as predicted by homology to Dprs and DIPs

We hypothesized that if IgLONs are homologous to Dprs and DIPs, and preserved their mode of action in the vertebrate line, they may similarly have homophilic and heterophilic binding activity. To test this, we employed the same high-throughput method, the extracellular interactome assay (ECIA), originally used in the discovery of Dpr-DIP interactions. In short, each ectodomain was cloned into two expression vectors, one in bait form using dimeric Fc fusions, and the other in prey form using pentameric Alkaline Phosphatase (AP_5_) fusions. The bait form can be captured on Protein A-coated plates through the high-affinity Fc-Protein A interaction, and capture of the prey by the bait is measured via AP activity using a chromogenic alkaline phosphatase substrate. The coating of the bait on plates and the highly oligomerized, parallel nature of the AP_5_ fusions mimic cell-to-cell adhesions, and increase apparent affinity via avidity by 10 to 1,000 fold (6).

In the binding assay, we also included the vertebrate Nectin and Nectin-like (Necl, or SynCAM for *synaptic cell adhesion molecule*) proteins. Like DIPs and IgLONs, Nectins and Necls have three Ig domains (13). They are known to interact homo- and heterophilically, mediate cell-to-cell interactions in the nervous and immune systems (13, 14), and form complexes structurally similar to Dpr-DIP complexes (8). When the ECIA was performed with all five mouse IgLONs and eight out of nine Nectins and Necls, we saw that Nectins and Necls interact with each other but do not produce complexes with IgLONs (Fig. 2B). All IgLONs interacted with each other, and could form both homophilic and heterophilic complexes. This is in agreement with our prediction based on the homology of IgLONs to Dprs and DIPs, and with previously reported interactions within the IgLON subfamily (15–18). While Nectins and Necls share many structural features with Dprs, DIPs and IgLONs, their lack of any interactions with IgLONs and the monophyly of Dprs, DIPs and IgLONs led us to conclude that they should be considered as a separate family within the IgSF (Figure 1).

Dprs, DIPs and IgLONs function through molecular interfaces to establish specific interactions on the cell surface. Next, we set out to determine whether Dprs, DIPs and IgLONs share functional interfaces and form complexes using a conserved surface. We hypothesized that if Dpr-DIP complexes and IgLON complexes are structurally related, structural models of IgLON complexes based on homology to Dprs and DIPs would accurately predict interface residues in IgLON complexes. Therefore, we created homology models of the IG1 domains of two IgLON complexes, one homophilic (OBCAM-OBCAM) and one heterophilic (NTM-NEGR1), based on the known structure of the Dpr6-DIP-α IG1-IG1 complex (8). Using these models, we have predicted four residues, labeled by asterisks in Fig. 2C and depicted on the NTM-NEGR1 model in Fig. 2D, to be at binding interfaces in IgLONs. When ECIA was repeated with single-site alanine mutants of these residues in OBCAM, NTM and NEGR1 against wild-type OBCAM, NEGR1 and NTM, respectively, we observed that two out of four mutants lost all detectable binding (Fig. 2E). The same positions in Dpr6 and DIP-α were previously identified to be crucial for the Dpr-DIP interaction (8). These results support a close evolutionary and structural relationship between IgLONs and Dprs and DIPs. Furthermore, mutations of the two hot spots of binding in IgLON complexes can be used as tools in future functional studies on IgLONs.

### Nematode homologs mimic binding activities of Drosophila Dprs and DIPs

We then focused on the predicted Dpr and DIP homologs of another protostome model organism, the nematode worm *Caenorhabditis elegans*. In *C. elegans*, we have identified one Dpr homolog, ZIG-8, and one DIP homolog, RIG-5 (Fig. 3A). Similar to Dprs and DIPs, ZIG-8 and RIG-5 have two and three Ig domains, respectively. They have an N-terminal signal peptide, and carry C-terminal hydrophobic sequences, indicative of membrane attachment via transmembrane helices (Fig. 3B). ZIG-8 was named as a member of the *z*wei-*Ig* class of proteins as it contains two Ig domains (19), and RIG-5 was assigned to the RIG class due to its being a neu*R*onal *Ig* protein. Interestingly, none of the other ZIG or RIG proteins belong to the Wirin family.

**Fig. 3.**
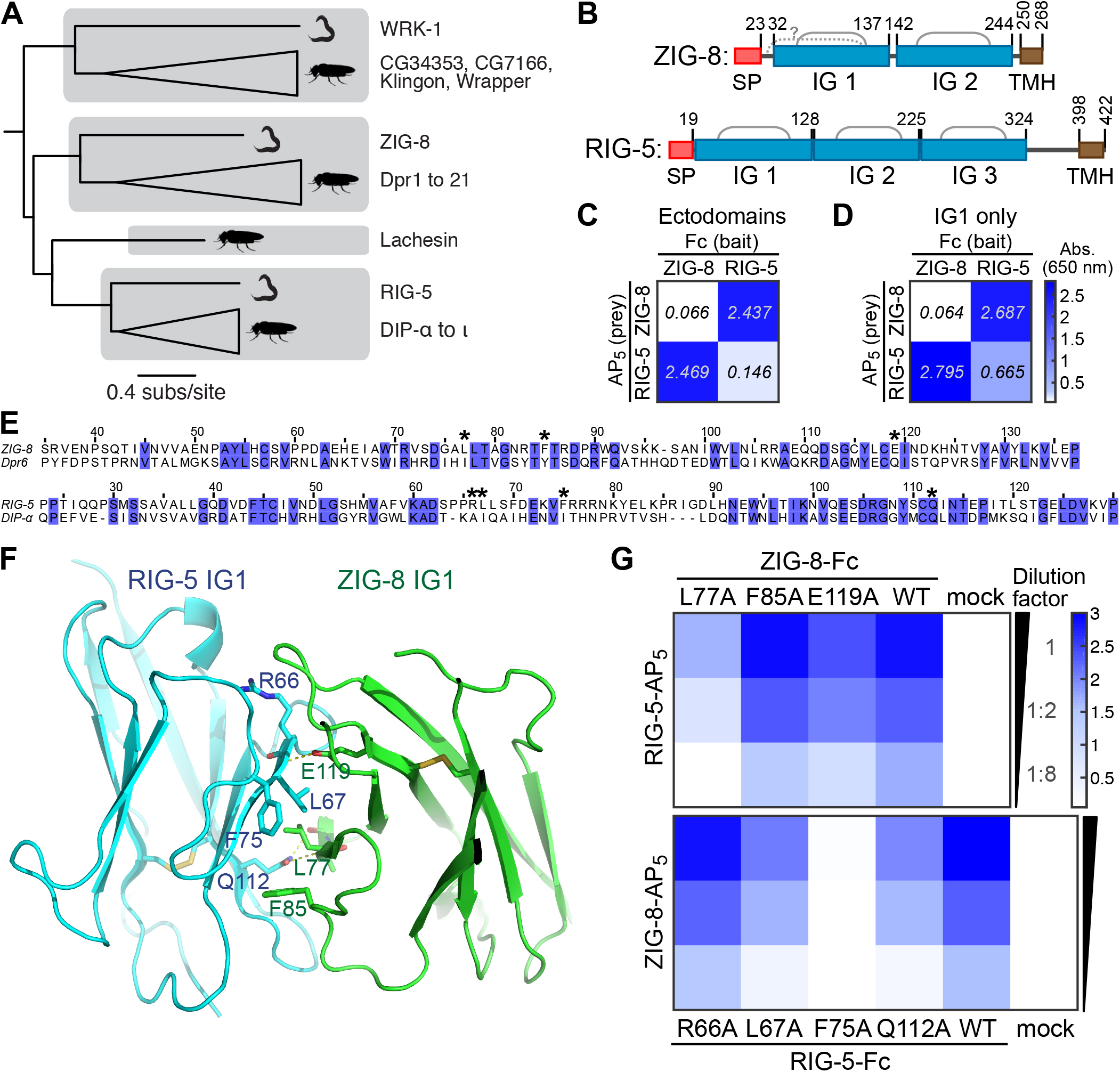
ZIG-8 and RIG-5, proposed Dpr and DIP orthologs in *C. elegans*, bind each other using interfaces identified in the *Drosophila* Dpr6-DIP-α complex structure. **A.** Phylogeny of Wirins in the two protostome model organisms, *C. elegans* and *D. melanogaster*. **B.** The domain structure of ZIG-8 and RIG-5. The gray arches represent predicted disulfides linkages. The question mark indicates an extra pair of cysteines that may form a disulfide bond. SP: Signal Peptide. TMH: Transmembrane helix. IG: Immunoglobulin domain. **C and D.** Binding experiments for ZIG-8 and RIG-5, performed using ECIA. The heterophilic interaction between ZIG-8 and RIG-5 is observed between both ectodomains (C) and IG1 domains only (D). RIG-5 forms a weaker IG1-IG1 homodimer, similar to DIPs in *Drosophila*. Fc (bait) and AP5 (prey) concentrations were normalized by dilutions within each panel using western blots to measure protein expression levels (see Fig. S4A). **E.** Sequence alignments of IG1 domains of ZIG-8 and Dpr6, and of RIG-5 and DIP-α. Asterisks indicate the four amino acid positions mutated in binding experiments. **F.** Homology model of ZIG-8-RIG-5 IG1-IG1 complex with mutated interface residues highlighted. E119 (ZIG-8) and Q112 (RIG-5) are predicted to make hydrogen bonds to main chain atoms on their interaction partners. **G.** ECIA for ZIG-8-RIG-5 heterodimer with mutations designed based on homology modeling. To effectively compare wild-type to mutants, protein concentrations within each mutant series were normalized. For each bait-prey pair, bait and prey were diluted two- and eight-fold to compare binding affinities at non-saturating concentrations.

The nematode *C. elegans* is an excellent candidate for the study of Dprs and DIPs, as it is a well-established model organism for neuronal wiring, and there are only one Dpr and one DIP in this species. Hence, we chose to characterize the two proteins and establish binding activity predicted based on their homology to *Drosophila* proteins. We performed ECIA with full-ectodomain constructs of ZIG-8 and RIG-5 (Fig. 3C), and found that they form a heterophilic complex. We also detected a weaker RIG-5 homodimer (Fig. 3C). As *Drosophila* Dprs and DIPs form heterophilic dimers (6), and DIPs can also form weaker homophilic dimers (10, 20), the binding activities of ZIG-8 and RIG-5 mimic those of their fly counterparts. As is the case for Dprs, DIPs and IgLONs, ZIG-8 and RIG-5 binding can be recapitulated only using the first IG domains (Fig. 3D).

Next, we showed that the *Drosophila* and *C. elegans* Dpr-DIP complexes are structurally similar, and that we can design binding mutants of ZIG-8 and RIG-5 that can be used in future genetic studies. Based on homology models, we identified the same four residues previously mutated in Dprs and DIPs (8, 10) and IgLONs (see Fig. 2) in ZIG-8 and RIG-5 (Figs. 3E-F). When these residues were mutated to alanine, the heterophilic binding between ZIG-8 and RIG-5 was weakened or broken (Fig. 3G). The most effective mutations for breaking the ZIG-8-RIG-5 complex were L77A (ZIG-8) and F75A (RIG-5) (see *Materials and Methods* for sequence numbering). In the homology model of the ZIG-8-RIG-5 complex, both residues are buried within holes on the interacting proteins’ surfaces (Fig. S4B-C), explaining why these residues are essential for ZIG-8-RIG-5 complex formation.

We also characterized the weak RIG-5 homodimer observed in ECIA. First, we investigated the solution properties of the RIG-5 IG1 domain, which was expressed, purified, and applied on a size-exclusion chromatography column at different quantities. Depending on its concentration, the elution volume of RIG-5 changed: when dilute, the elution volume for RIG-5 was close to that expected for a monomer, but when concentrated, RIG-5 ran like a dimer (Fig. 4A). This behavior indicates a fast-exchange monomer-to-dimer equilibrium at concentrations tested. Overall, we observed that RIG-5 IG1 can form homodimers with a dissociation constant in the mid-to-high micromolar range (Fig. 4B). These results closely mimic size-exclusion behavior of the homodimeric *Drosophila* DIP-α and DIP-η (10).

**Fig. 4.**
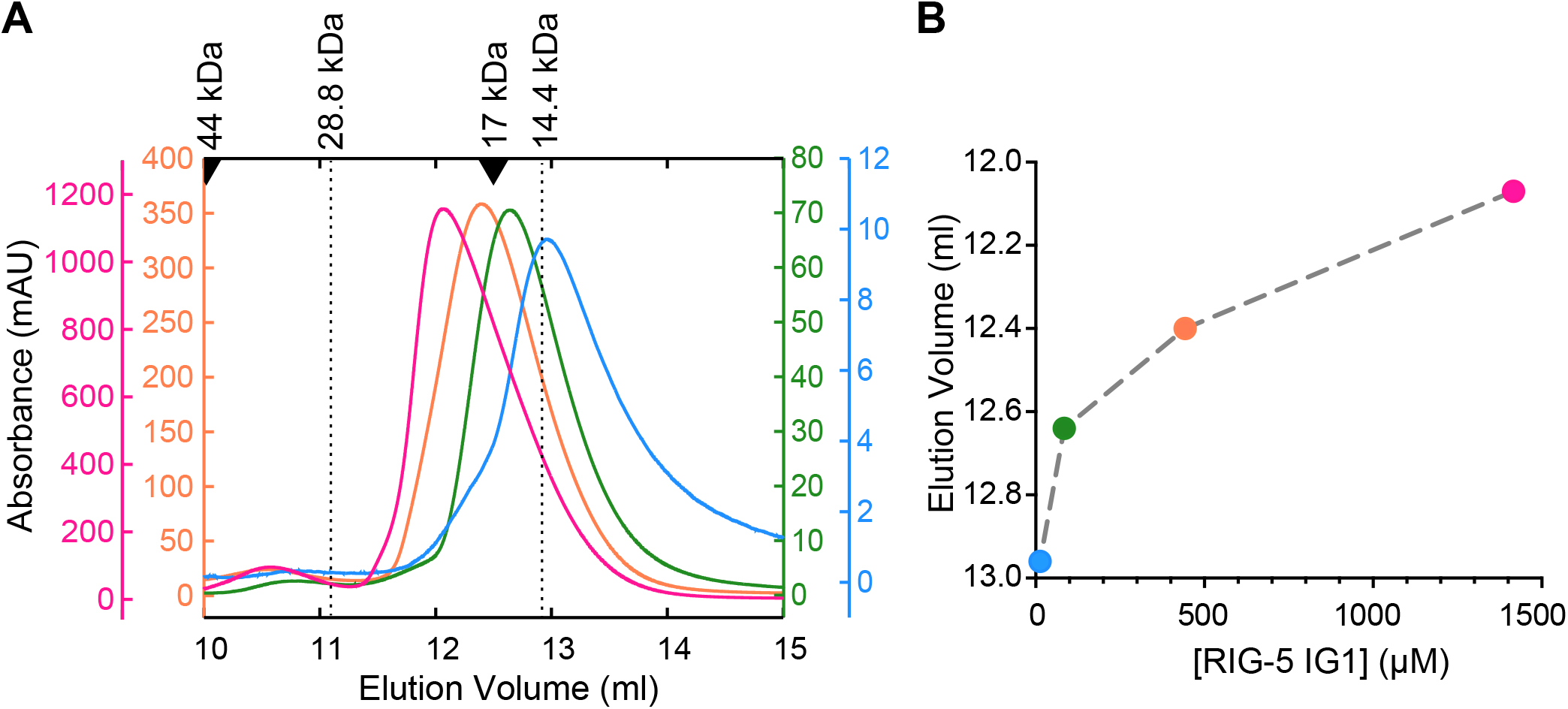
The IG1 domain of the *C. elegans* DIP ortholog RIG-5 is a weak dimer in solution, similar to *Drosophila* DIP-α and -η. **A.** RIG-5 IG1 elution volumes in size-exclusion chromatography decreases with increasing RIG-5 concentration in the μM range, indicative of a fast-exchange monomer-dimer equilibrium. Elution volumes for molecular weight standards are shown above as arrowheads. The RIG-5 IG1 construct used encodes for a 13.2 kDa protein product, which yields a ~14.4 kDa mature glycoprotein. **B.** Elution volumes plotted against loaded protein concentration indicates an dissociation constant in the mid-to-high micromolar range for the RIG-5 homodimer, similar to what was observed for DIP-α and DIP-η (10).

Finally, some of the common features of Dprs and DIPs are the relative flexibility between the IG domains, stretches of unstructured and/or low-complexity regions in the juxtamembrane regions, and the lack of intracellular domains. These observations support the idea that Dprs and DIPs do not signal directly, but are adhesive receptors that utilize yet-to-be-identified co-receptors or secreted ligands for signaling for neural wiring processes through pathways such as the retrograde BMP signaling pathway (8). Overall, RIG-5 and ZIG-8 share these sequence features (Fig. 3B).

### The crystal structure of the RIG-5 homodimer

Many IgSF proteins are known to homodimerize, as has been observed here for Nectins, IgLONs and DIPs, and do so commonly in the micromolar range. To demonstrate that the weak RIG-5 homodimers we observed are structurally similar to DIP homodimers, we determined the crystal structure of the first immunoglobulin domain of RIG-5 to 1.42 Å resolution (Table 1 and Fig. 5A). RIG-5 IG1 can be superimposed on Dpr or DIP IG1 domains solved to-date, with root-mean-square deviation (rmsd) of Cα atoms at 0.9-1.0 Å, indicative of homology between RIG-5 and Dprs and DIPs (Fig. 5B). The structure shows that RIG-5 forms symmetrical homodimers using one of its β-sheets including the *GFCC’C”* strands, and the overall geometry of interaction within the RIG-5 homodimer closely resembles binding in the *Drosophila* Dpr-DIP and DIP-DIP complexes (Fig. 5B). The rmsd of the RIG-5 complex from five experimentally determined Dpr-DIP and DIP-DIP structures is 2.62 Å ± 0.12 Å (standard deviation). The interface area for the RIG-5 dimer is 940 Å^2^, slightly above the interface areas observed for Dpr and DIP complexes, which range from 820 to 910 Å^2^ (excluding glycans) (8, 10).

**Table 1.**
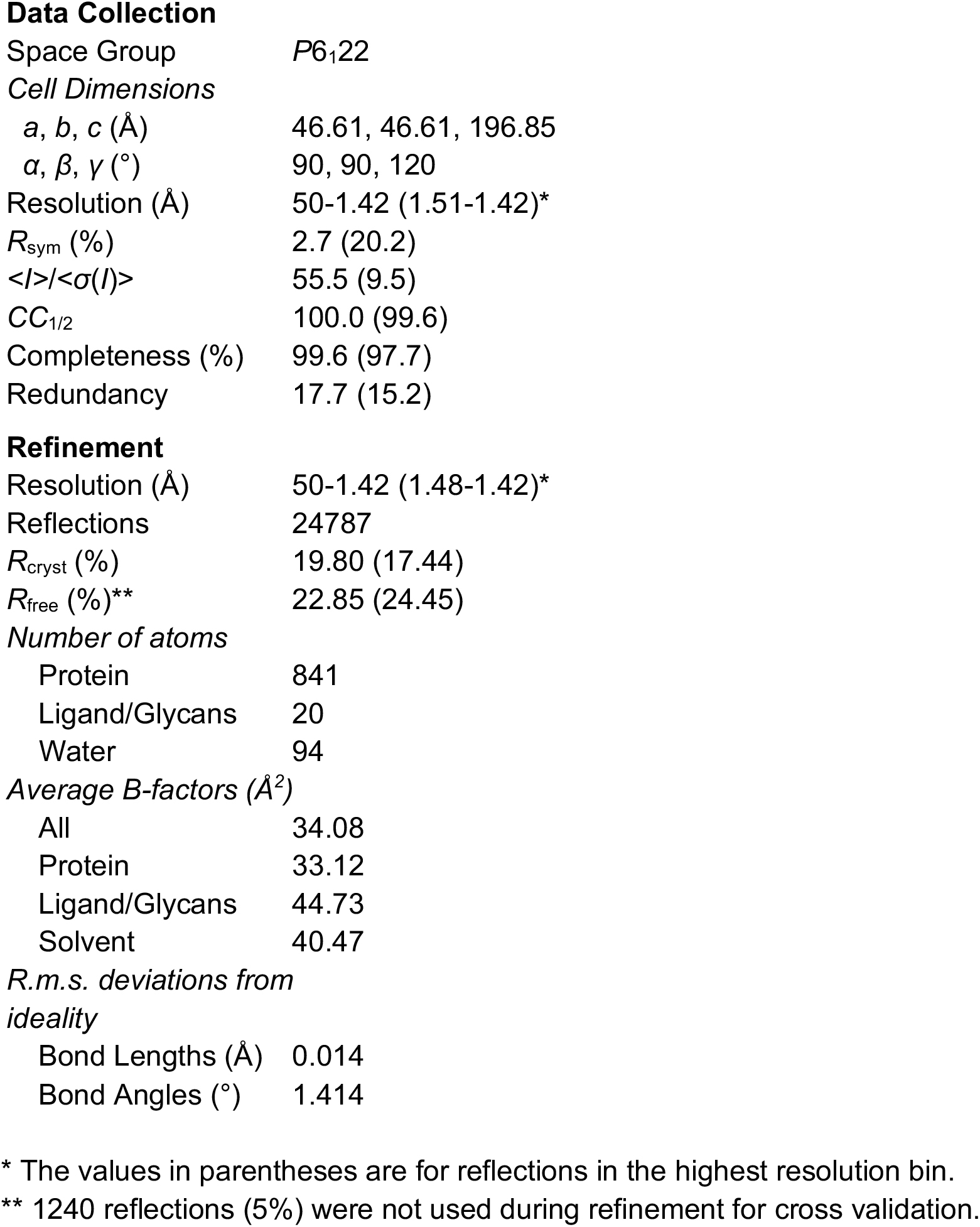
Data and refinement statistics for x-ray crystallography of the RIG-5 homodimer.

|  |  |
| --- | --- |
| Space Group | *P*6<sub>1</sub>22 |
| *Cell Dimensions* |  |
| *a*, *b*, *c* (Å) | 46.61, 46.61, 196.85 |
| $\alpha$, $\beta$, $\gamma$ (°) | 90, 90, 120 |
| Resolution (Å) | 50-1.42 (1.51-1.42)* |
| *R*<sub>sym</sub> (%) | 2.7 (20.2) |
| $\langle I \rangle / \langle \sigma(I) \rangle$ | 55.5 (9.5) |
| *CC*<sub>1/2</sub> | 100.0 (99.6) |
| Completeness (%) | 99.6 (97.7) |
| Redundancy | 17.7 (15.2) |
| **Refinement** |  |
| Resolution (Å) | 50-1.42 (1.48-1.42)* |
| Reflections | 24787 |
| *R*<sub>cryst</sub> (%) | 19.80 (17.44) |
| *R*<sub>free</sub> (%)** | 22.85 (24.45) |
| *Number of atoms* |  |
| Protein | 841 |
| Ligand/Glycans | 20 |
| Water | 94 |
| *Average B-factors (Å<sup>2</sup>)* |  |
| All | 34.08 |
| Protein | 33.12 |
| Ligand/Glycans | 44.73 |
| Solvent | 40.47 |
| Bond Lengths (Å) | 0.014 |
| Bond Angles (°) | 1.414 |

\* The values in parentheses are for reflections in the highest resolution bin.
\*\* 1240 reflections (5%) were not used during refinement for cross validation.

**Fig. 5.**
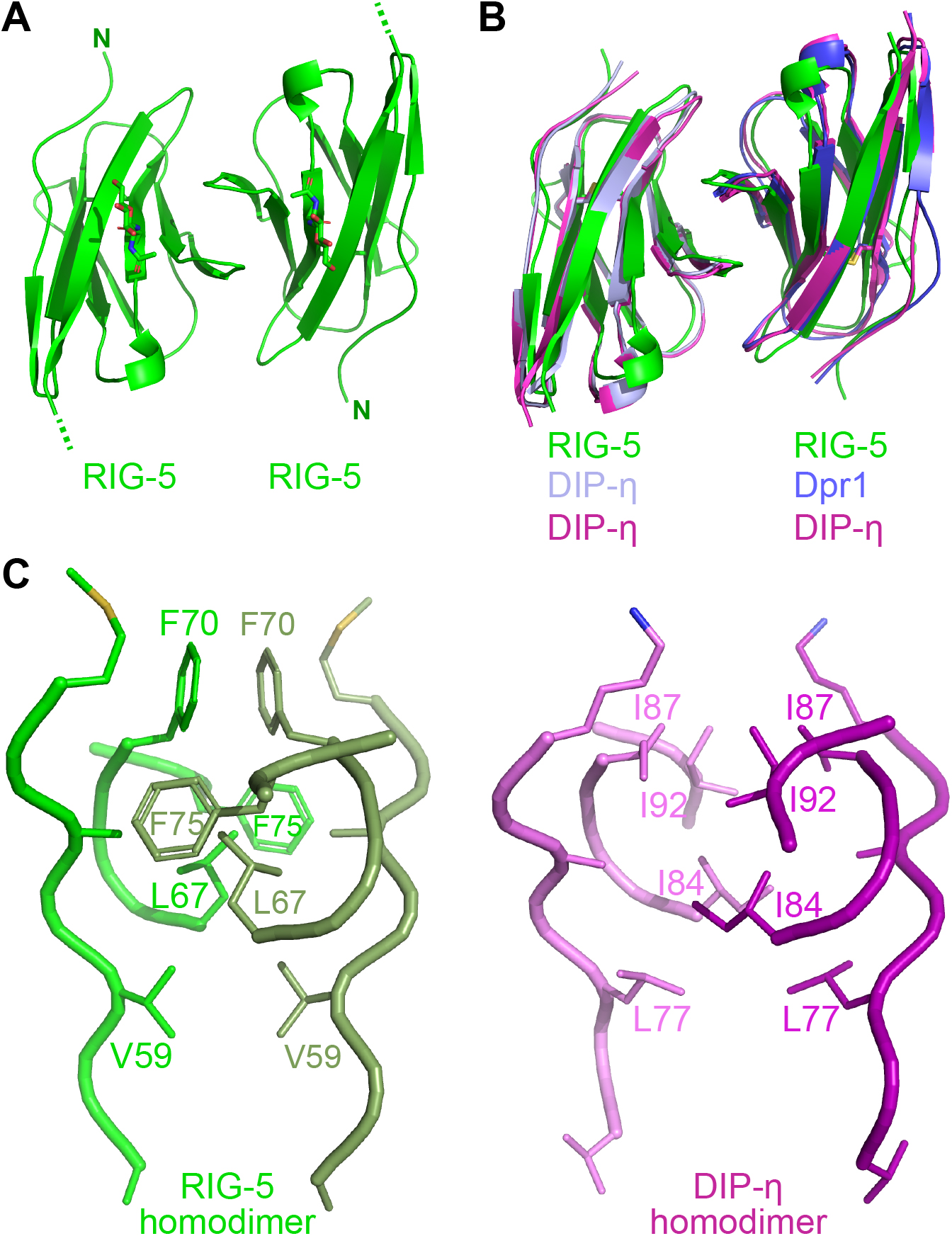
Crystal structure of the RIG-5 homodimer highlights homology with Drosophila DIPs. **A.** Crystal structure of the RIG-5 IG1 homodimer. One N-linked glycan per monomer is depicted in sticks representation. **B.** Overlaid structures of the RIG-5 and DIP-η homodimers, and the Dpr1-DIP-η heterodimer. **C.** Comparison of homodimeric interfaces in RIG-5 and DIP-η structures.

The detailed chemical features of the RIG-5 dimerization interface are highly similar to those of the Dpr and DIP interfaces (Fig 5C). A central hydrophobic core is surrounded by permissive polar interactions (Fig. S5). A comparison of the hydrophobic core residues in the RIG-5 and DIP-η homodimeric structures show that while every amino acid is different at the core, the differences are conservative: for example RIG-5 L67 is equivalent to DIP-η I84 (Fig. 5C).

Furthermore, despite complete lack of sequence identity at the interface cores, the main chain positions at the RIG-5 and DIP homodimer interfaces are highly similar (Figure 5C). The RIG-5 homodimeric interface is characterized by high shape complementarity (*sc* = 0.73) (21), probably to make up for lack of charge complementarity, as we previously observed also for Dpr-DIP and DIP-DIP structures (8, 10).

## DISCUSSION

Immunoglobulin superfamily proteins went through a large expansion in the metazoan line to support many multicellular functions, including roles in establishing neuronal connectivity. IgSF proteins have been implicated in a multitude of wiring functions, including neurite outgrowth and axon guidance, axon coalescence and fasciculation into bundles, and synaptic morphogenesis and targeting (22). We and others have identified many interactions among IgSF proteins that are likely to mediate cell-cell interactions involved in these functions. However, much remains to be discovered; even after the completion of large-scale IgSF interactome screens, a large fraction of IgSF proteins remain as orphans (i.e., their binding partners are unknown) (6).

### Members of the Wirin family perform conserved functions

The evolutionary, biochemical and structural relationship we have established between Dprs and DIPs in flies and IgLONs in vertebrates are further corroborated by a collection of functional similarities. As discussed above, Dprs and DIPs are expressed in small subsets of neurons and regulate retrograde signaling, synaptic targeting, and cell death. The expression and localization of IgLONS have been studied in the developing and adult rodent brain. Members of the subfamily have different spatial and temporal expression patterns (14, reviewed in 23). IgLONs, are expressed both in neurons and oligodendrocytes (24), and have been observed to be localized to pre- and/or post-synaptic membranes in developing and adult brains, lending support for a synapse formation and maintenance function. NTM is known to localize to granule cell-Purkinje cell and mossy fiber-granule cell synapses in the cerebellum (25), while LSAMP is observed postsynaptically in granule cells of the dentate gyrus in the adult mouse hippocampus (26), and NEGR1 and OBCAM are found in post-synaptic densities in the cerebral cortex and the hippocampal CA3 region in rats (27). Synaptic localization is further validated by a recent proteomic analysis of synaptic clefts, where the 199-member proteome of excitatory synapses in rat embryo cortical neurons included all five members of the IgLON subfamily. Interestingly, NEGR1, OBCAM, NTM and LSAMP were also identified among proteins of the synaptic vesicle proteome in rat brains (28). In addition, support for neurite growth or axon fasciculation functions for several IgLONs have been reported (for example, see 25). These data suggest that the vertebrate and arthropod members of the Wirin family, IgLONs and Dprs/DIPs, share functional roles in establishing connectivity in their respective nervous systems.

Our phylogenetic analysis has highlighted additional Wirin subfamilies in protostomes: Klingons and Lachesins. Several observations also connect these proteins to Dprs, DIPs and IgLONs. Most strikingly, *Drosophila* Klingon interacts with a secreted leucine-rich repeat (LRR) domain protein called cDIP (common Dpr and DIP interacting protein), which was named for its ability to interact with most Dprs and DIPs (6). Klingon is necessary for long-term memory formation (29) and is involved in the development of the fly photoreceptor neuron R7 (30). It is not known if Klingon and Dpr11, which is selectively expressed in one subclass of R7 neurons (8), cooperate in R7 development or connectivity. Also, similarities between *Drosophila* Lachesin and vertebrate IgLONs, both containing three IG domains and a GPI anchor for membrane attachment, were previously recognized due to shared domain features (31, 32). Lachesin is expressed in neuronal (31), epithelial (33) and glial populations (32); its function remains poorly understood.

Finally, we have identified the nematode orthologs of the Wirin family. ZIG-8, the Dpr ortholog, was first identified as one of the candidate ZIG genes involved in the maintenance axons within the ventral nerve cord in *C. elegans* (19, 34). RIG-5, the DIP ortholog, has been implicated in the navigation of axons within the ventral nerve cord (35). Among protostomes, given the relative simplicity of its nervous system, it is intriguing to speculate that *C. elegans* has a limited Dpr/DIP repertoire (two genes), while the complex nervous systems of arthropods and mollusks utilize expanded sets of Dpr/DIP genes (thirty in the fruit fly).

### A wider neuronal wiring family includes Kirrels, Nephrins, Nectins and Necl/SynCAMs

In our attempts to find proper outgroups for creating a phylogenetic tree for Wirins, we have also recognized that four other protein families, Kirrels, Nephrins, Nectins and Necls, are distantly related to Wirins. We have previously reported (8) remarkable structural similarities among the structures of the Dpr6-DIP-α complex, the Kirrel-Nephrin complexes (36), and known Nectin and Necl homo- and heterophilic complexes (37–40). Recent work has also identified functional similarities between Kirrels and Dpr/DIPs in the organization of olfactory sensory neurons in mammals and flies (12). Wirins and these proteins interact with their homo and heterophilic partners using the *GFCC’C”* faces of their N-terminal IG domains, which is a feature, to our knowledge, not observed in other neuronal IG complexes (Fig. S6). Furthermore, Wirins and the four distantly related families of proteins appear to adopt structures with fully extended ectodomains, unlike other neuronal families, such as DSCAMs, DCC and Axonin, where IG domains fold back to create horseshoe-shaped structures. Finally, Kirrels, Nephrins, Nectins and Necl/SynCAMs are well-studied molecules with recognized functions in establishing neuronal connectivity.

The structural similarities between Wirins and the four distantly related families, and the topological similarities between their complexes break down beyond the first three immunoglobulin domains. Kirrels and Nephrins contain five and ten extracellular domains, respectively, unlike the two and three domains observed in most Wirins, Nectins and Necls. Kirrels, Nephrins, Nectins and Necls contain conserved intracellular regions specialized for signaling, while the Wirins usually do not have intracellular domains as they are anchored to the plasma membrane with GPI linkages or via transmembrane helices located at their extreme C termini. There are exceptions to this among Dprs, in that isoforms of some Dprs have sequences that are likely to encode intracellular domains. However, these putative intracellular domain sequences are poorly conserved between species, and some of these isoforms may not be expressed.

### A shared structural architecture in neuronal IgSF proteins

These connections established and cited between the Dpr, DIP and IgLON families, Klingon, Lachesin, and their nematode orthologs help define a functional family of proteins with a shared structural architecture involved in the establishment of neuronal connectivity going back at least to the rise of bilaterians. Future studies will investigate the evolutionary origins of the wider family of neuronal wiring molecules, which would require further identification of Wirin-like proteins in Cnidaria. It would be of interest to see if the Wirins and its four related IgSF protein families arose during the period in which neurons and neuronal circuits first appeared. Finally, expansion of the Wirin family appears to correlate with the evolution of nervous systems of increasing complexity especially in protostomes.

## MATERIALS AND METHODS

### Phylogenetics

Putative Dpr and DIP homologs were identified using BLAST with *D. melanogaster* proteins as queries (41). To exclude distant IgSF homologs, the BLAST hits were used as queries to search over the *D. melanogaster* proteome (reciprocal BLAST), and only those with a Dpr or DIP as the top hit were retained. For identifying IgLON homologs in protostomes, human IgLONs were used as queries and reciprocal BLAST was performed on the human proteome. Amalgam, CG34353, CG7166, DIP, Dpr, Klingon, Lachesin, and Wrapper were identified as IgLON homologs in *D. melanogaster*. Only some of the 21 Dprs appeared as IgLON homologs, presumably due to their fast rate of evolution. Amalgam was excluded from analysis because it could not be reliably aligned to other proteins. Nectin, Necl, Kirrel, and Nephrin were the only other IgSF subfamilies that could be reliably aligned to the Wirin family.

Sequences for insect, bony fish, and tetrapod proteins were obtained from the NCBI protein database. To acquire sequences from other organisms, whose proteomes were generally poorly represented in the NCBI protein database, transcripts were assembled from RNA-seq reads in the NCBI SRA database (42). To selectively assemble specific transcripts rather than the entire transcriptome, RNA-seq reads with similarity to reference proteins were extracted using TBLASTN. Transcripts were then *de novo* assembled using Velvet/Oases (43, 44), and coding sequences were identified using TransDecoder (http://transdecoder.github.io) (45).

Only the Ig domains were included in the alignment because the rest of the proteins could not be aligned across paralogs. The number of Ig domains varied across proteins: two for Dpr, three for DIP, IgLON, Klingon, Lachesin, Nectin, and Necl, five for Kirrel, and ten for Nephrin. The first one and a half Ig domains of Dpr aligned with the corresponding part of the 3-Ig proteins. The rest of Dpr aligned with the second half of the third Ig domain of the 3-Ig proteins. The first two Ig domains of the 3-Ig proteins aligned with the corresponding part of Kirrel and Nephrin. The alignment of the third Ig domain to the rest of Kirrel and Nephrin was ambiguous. However, the phylogeny was robust to the uncertainty in alignment. When alignments were generated using different gap penalties, the resulting phylogenies were topologically identical to that in Fig. 1. Furthermore, using only the first two Ig domains resulted in a phylogeny that is also topologically identical to that in Fig. 1.

Based on preliminary alignments and phylogenies, proteins with exceptionally long branches or nonsensical species placements were discarded. The remaining proteins were classified into paralog groups. Proteins in each paralog group were aligned separately, removing paralog-specific insertions and alignment-ambiguous regions. The full alignment was assembled from paralog alignments through sequential profile alignment. All alignments were generated using MUSCLE with default settings (46). Preliminary phylogenies were inferred using FastTree2 (47). The ML phylogenies were inferred using RAxML v8.2.12 (48). The best-fit model of evolution for the full phylogeny was WAG + G + I + X (X: ML estimation of equilibrium amino acid frequencies). The best-fit model for DIP, IgLON, and Klingon individually, however, were LG + G + I + X. Therefore, paralog relationships within each family were inferred in separate analyses.

Approximate likelihood ratio statistics were calculated using PhyML v.3.0 under the topology and equilibrium amino acid frequencies inferred using RAxML (49).

### The Extracellular Interactome Assay (ECIA)

Interactions between ectodomains were tested using ECIA (6) with minor modifications: The promoters in the bait and prey expression vectors have been replaced with the constitutively active Actin 5C promoter from *D. melanogaster* in lieu of the inducible metallothionein promoter, which improves expression levels, and therefore the sensitivity of the assay (unpublished results). The transfection agent was also changed to TransIT-Insect (Mirus), which was used according to the manufacturer’s recommended protocol. Mouse IgLON cDNAs were used in the binding experiments.

### Homology modeling

Homology modeling of Dpr and DIP orthologs using the Dpr6-DIP-α structure (PDB: 5EO9) was done with MODELLER (50). Further side chain rotamer optimization was performed using SCWRL4 (51) and manual inspection of alternate rotamers in PyMOL (52). For the NEGR1-NTM IG1-IG1 complex, Dpr6 was used to model NEGR1, DIP-α and was used to model NTM.

### RIG-5 sequence numbering

In the manuscript, we employed a numbering scheme for RIG-5 that used the C36F7.4g.1 transcript starting at the second methionine of the annotated transcript, resulting in 60 amino acids removed from the N terminus. See *supplemental information* for details.

### Expression and Purification of RIG-5 IG1

The N-terminal domain *C. elegans* RIG-5 was cloned into pAcGP67A with a C-terminal hexahistidine tag, and co-transfected into Sf9 cells with linearized baculoviral DNA (Expression Systems) using the TransIT-Insect transfection reagent (Mirus). Amplified virus was used to infect High Five cells. Media was collected 60 hours post-infection. RIG-5 was first purified using Ni-NTA agarose resin, followed by size-exclusion chromatography on a Superdex 75 10/300 column (GE Healthcare) in 10 mM HEPES, pH 7.2 and 150 mM NaCl.

### Crystallography of RIG-5 IG1

RIG-5 IG1 sample was concentrated to 17 mg/ml, and crystallized by the sitting-drop vapor diffusion method using a Mosquito crystallization robot (TTP Labtech) with 100 nl protein + 100 nl crystallant drops against a 50-μl crystallant reservoir. Best crystals grew in 0.1 M HEPES pH 7.0 and 1 M sodium citrate. Crystals were cryoprotected in 0.1 M HEPES pH 7.0, 1.2 M sodium citrate, 10% glycerol and vitrified in liquid nitrogen. Diffraction data were collected at SSRL beamline 9-2. Structure was solved by molecular replacement method using a DIP-η IG1 monomer (10) as the model in *PHASER* (53). The model was further refined with *phenix.refine* (54) and real-space model building was performed in *Coot* (55). Model validation was performed using *Molprobity* (56) within the *PHENIX* suite (57).

## ACKNOWLEDGMENTS

We thank Yeonhee Jenny Park for technical help, and Paschalis Kratsios for discussions. This work was supported in part by National Institutes of Health Grant R01 NS097161 (to E. Ö.), R37 NS028182 (to K. Z.), and R01 NS096509 (to K. Z.), and a Klingenstein-Simons Fellowship Award in the Neurosciences (to E. Ö.). Use of the Stanford Synchrotron Radiation Lightsource, SLAC National Accelerator Laboratory, is supported by the U.S. Department of Energy, Office of Science, Office of Basic Energy Sciences under Contract No. DE-AC02-76SF00515. The SSRL Structural Molecular Biology Program is supported by the DOE Office of Biological and Environmental Research, and by the National Institutes of Health, National Institute of General Medical Sciences (including P41GM103393).

## SUPPLEMENTARY INFORMATION

### Sequence numbering for RIG-5

There is ambiguity with regards to the N-terminal end of RIG-5 covering the signal peptide. The C36F7.4f.1 transcript on the Wormbase database (1) only allows for a weak prediction of a signal peptide with Phobius or SignalP (2, 3), while the C36F7.4g.1 transcript yields a strongly predicted signal peptide (“MYLFALLCGVLLVFKQACSRG”) if the second methionine in the transcript is used as the start methionine and therefore sixty amino acids are removed from the transcript. We used a numbering scheme throughout the manuscript that uses the C36F7.4g.1 sequence with 60 amino acids removed from the N terminus. It should be noted that the mature proteins (i.e. after the signal peptides are processed) for both transcripts have identical sequences. In the manuscript, we do not make a call about which transcript(s) are actually expressed in worms.

### Definitions for protein families within the IgSF

Nectins and Necls are two related families within the IgSF found in vertebrates. In humans, the family include nine members, Nectins 1 to 5 and Necls 1 to 4. It should be noted that Necl5 is more closely related to Nectins than Necls.

Here, we define Kirrels as orthologs of the *C. elegans* protein SYG-1. Kirrels are found across bilaterians, including SYG-1, Rst and Kirre (Duf) in *Drosophila*, and Kirrel1 (Neph1), Kirrel2 (Neph3) and Kirrel3 (Neph2) in vertebrates.

Nephrins are heterophilic binding partners of Kirrels, and are orthologs of the *C. elegans* protein SYG-2. Nephrins are found across bilaterians, including SYG-2, SNS and Hibris in *Drosophila*, and Nephrin in vertebrates.

### Constructs used in ECIA experiments

For ECIA experiments, expression constructs included the following residues.

ZIG-8 full-length ectodomain: Ala22 to Ser249 (Wormbase CDS Y39E4B.8).

ZIG-8 IG1: Ala22 to Pro137.

RIG-5 full-length ectodomain: Arg20 to Arg397 (Wormbase CDS C36F7.4e minus the N-terminal 60 amino acids).

RIG-5 IG1: Arg20 to Pro130.

Mouse OBCAM full-length ectodomain: Thr30 to Asn14 (NCBI Accession NP_808574.2)

Mouse NTM full-length ectodomain: Gly34 to Asn321 (NCBI Accession NP_758494.2)

Mouse NEGR1 full-length ectodomain: Val32 to Gly318 (NCBI Accession NP_ 001034183.1).

**Fig. S1.**
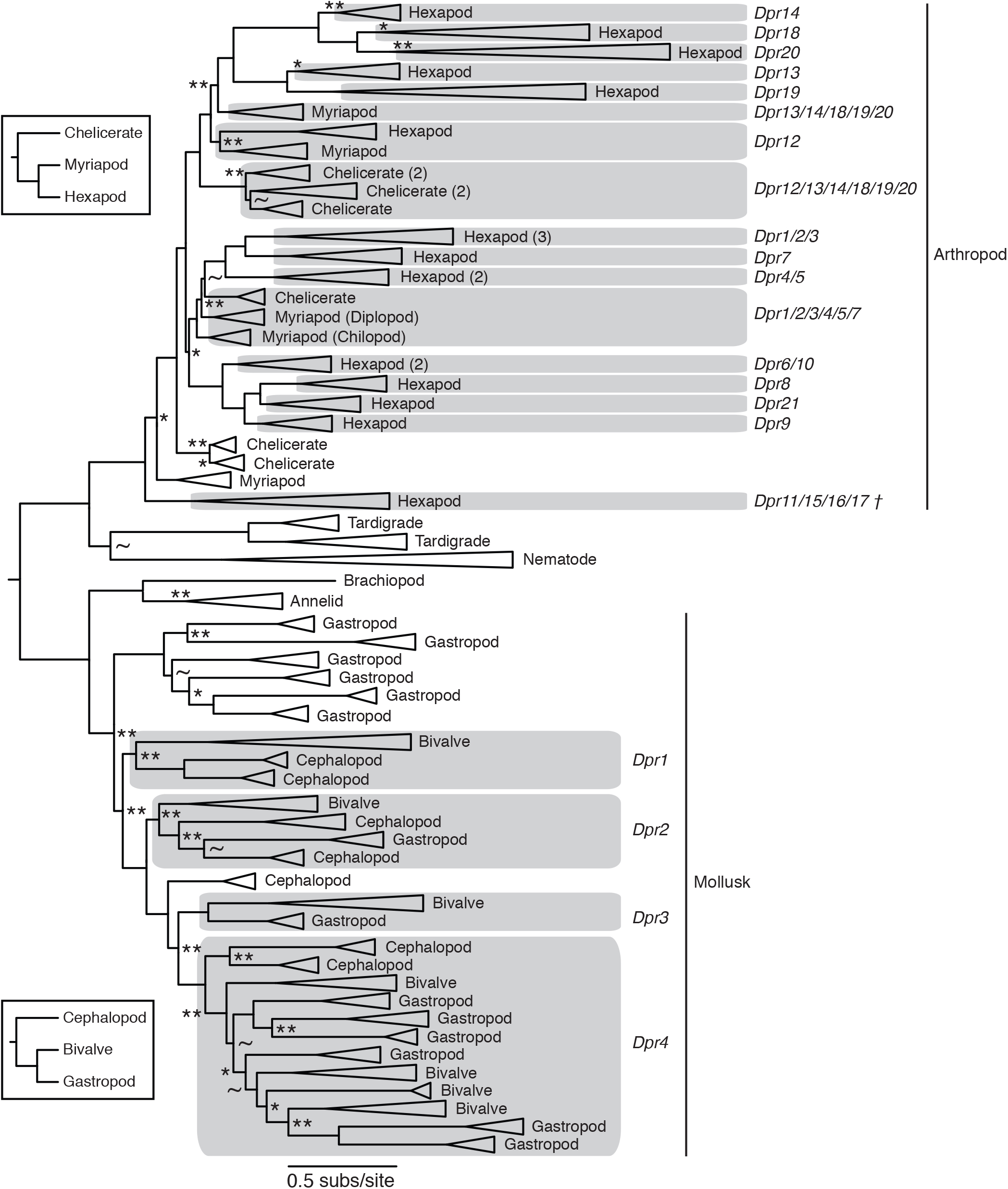
The ML phylogeny of the Dpr family. Approximate likelihood ratio statistics (aLRS) are shown as branch supports: ** < 9.2 (= 2 ln100), * < 4.6 (= 2 ln10), ~ < 2.2 (= 2 ln3). Unmarked branches have aLRS > 9.2. The insects show the arthropod and mollusk phylogenies. The arthropod paralogs are labeled following the D. melanogaster Dpr nomenclature. The numbers next to some clades show the maximum number of paralogs in the clade when there are gene duplications in its subclades.

**Fig. S2.**
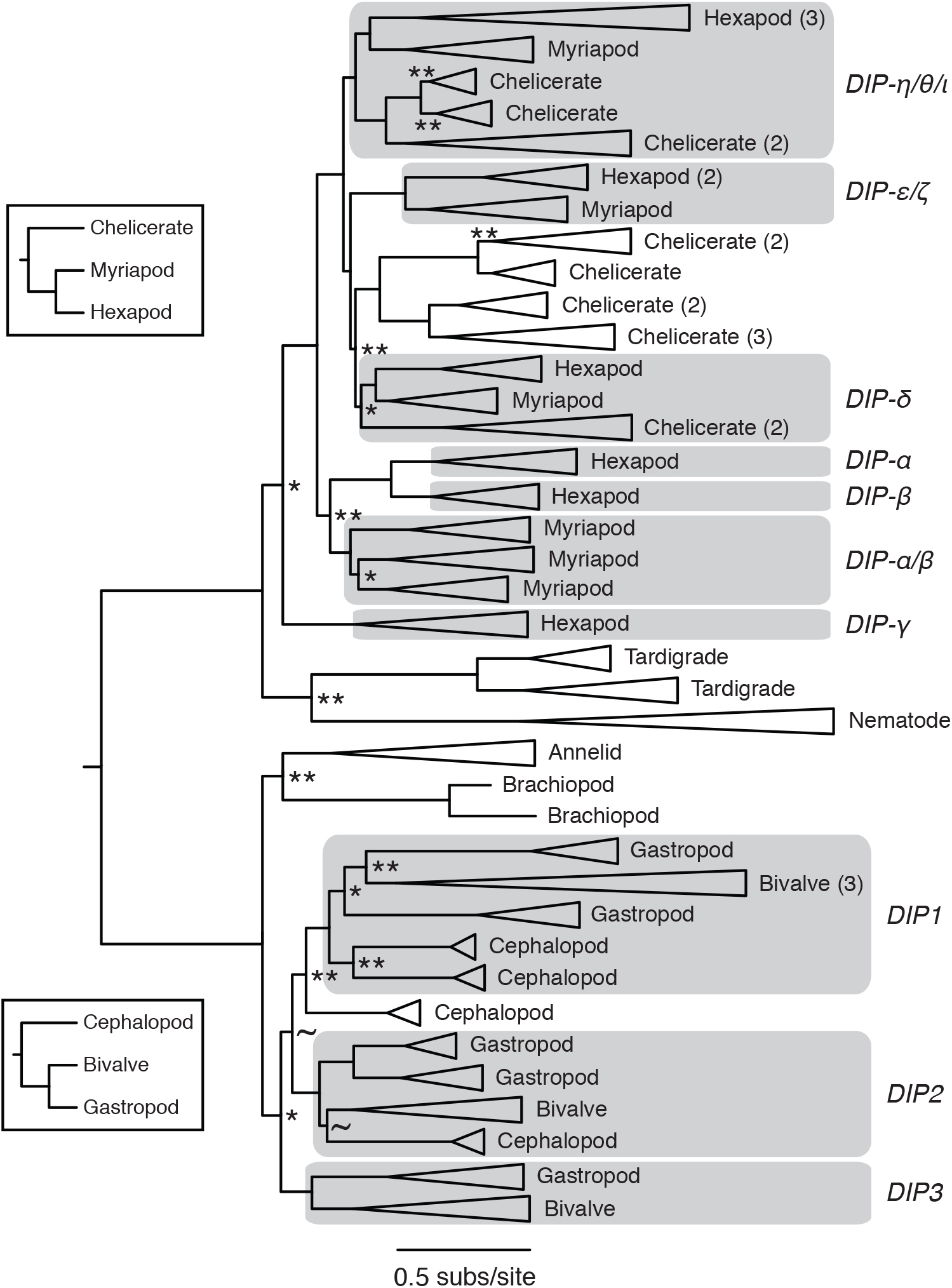
The ML phylogeny of the DIP family shown as in Fig. S1.

**Fig. S3.**
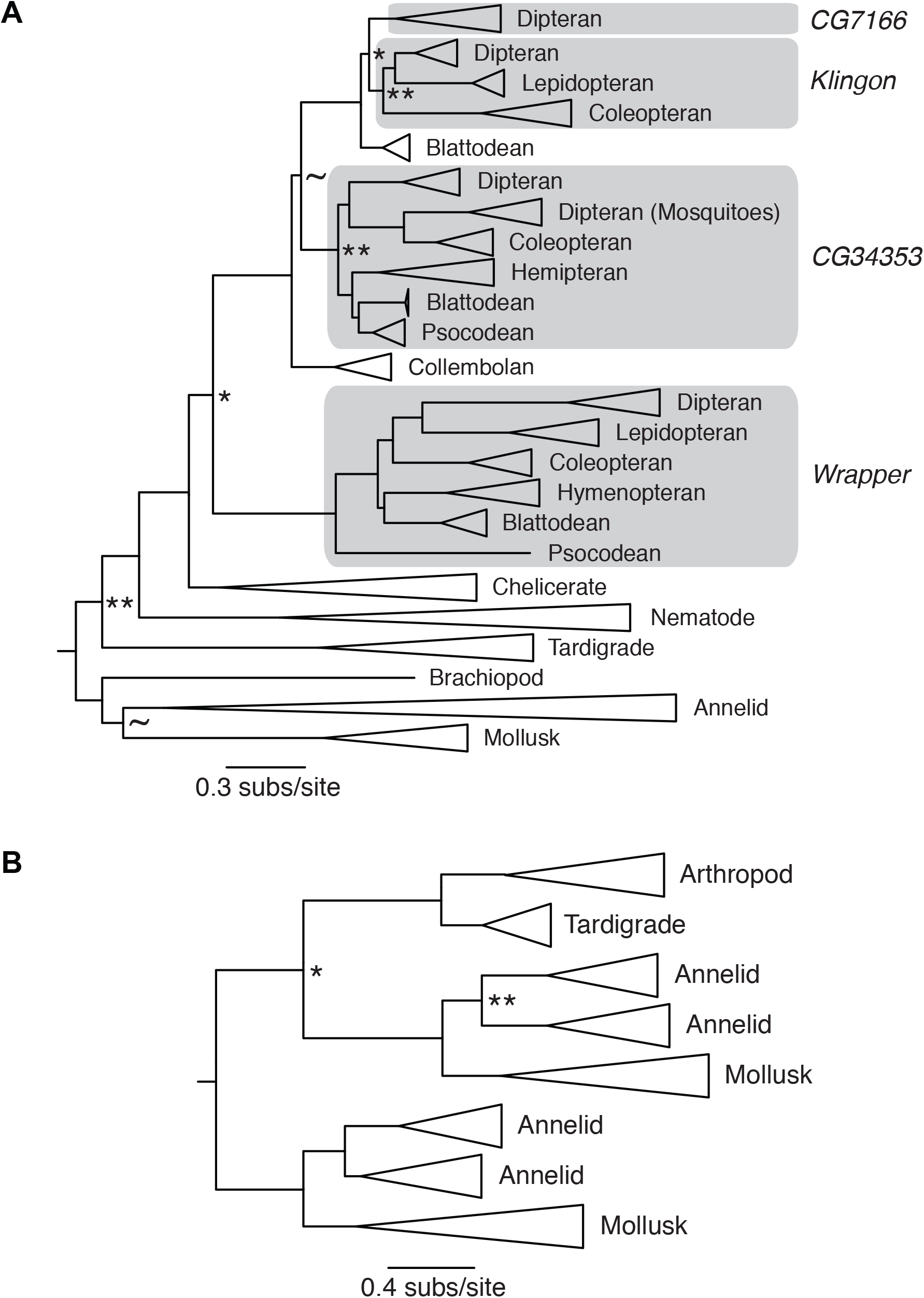
**A.** The ML phylogeny of the Klingon family shown as in Fig. S1. **B.** The ML phylogeny of the Lachesin family shown as in Fig. S1.

**Fig. S4.**
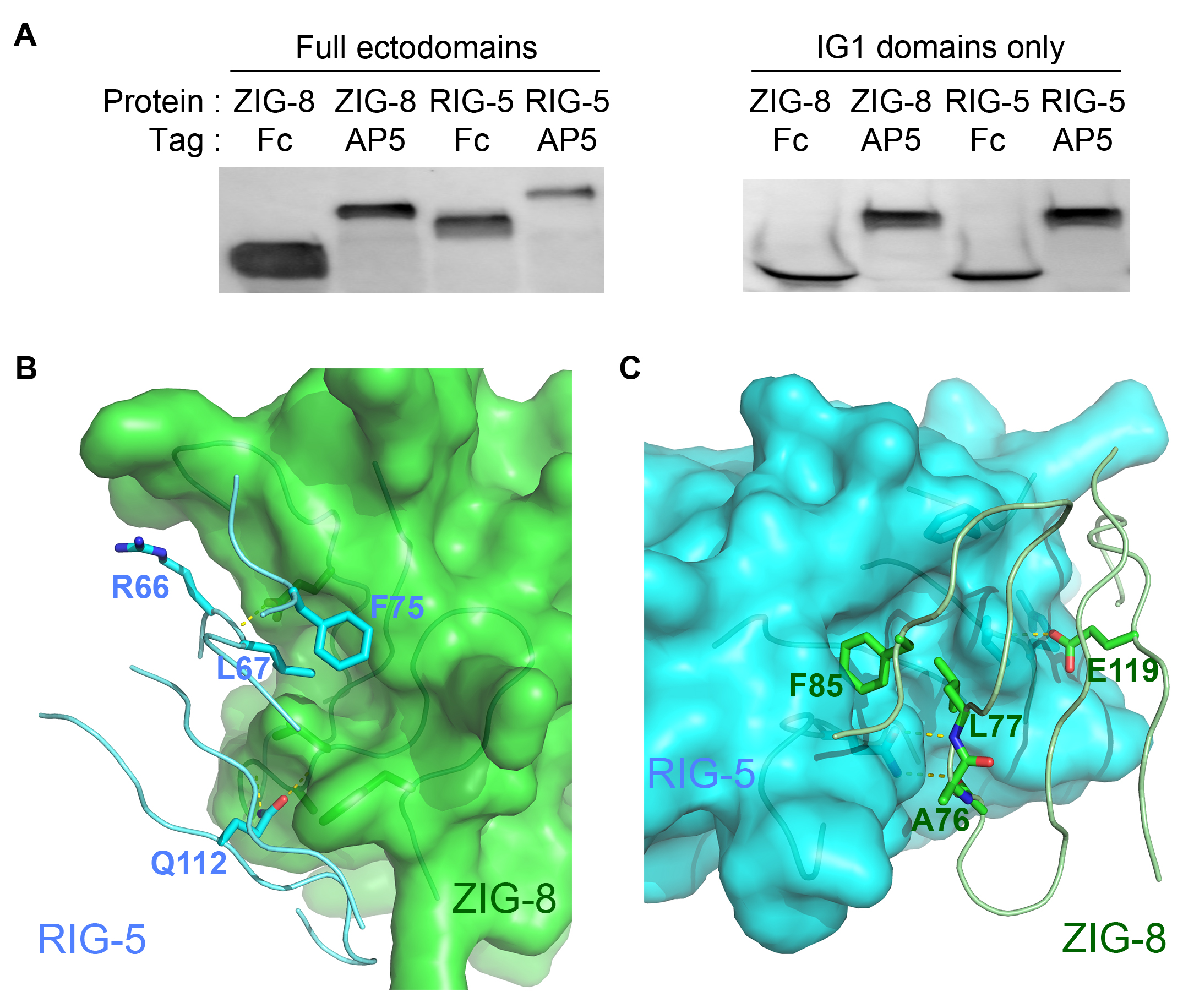
ZIG-8 and RIG-5 are predicted to bind each other through complementary surfaces. **A.** Western blots of conditioned media for ZIG-8 and RIG-5 proteins used in Fig. 3B and 3C. These blots were used to normalize protein concentrations for the ECIA. **B and C.** ZIG-8 (green) and RIG-5 (cyan) surfaces can accommodate hydrophobic side chains of F75 and L67 (RIG-5) and F85 and L77 (ZIG-8), based on homology modeling. All side chains mutated in Fig. 3F are displayed as sticks, as well as the main chain atoms predicted to make hydrogen **bonds to Q**112 (RIG-5) and E119 (ZIG-8).

**Fig. S5.**
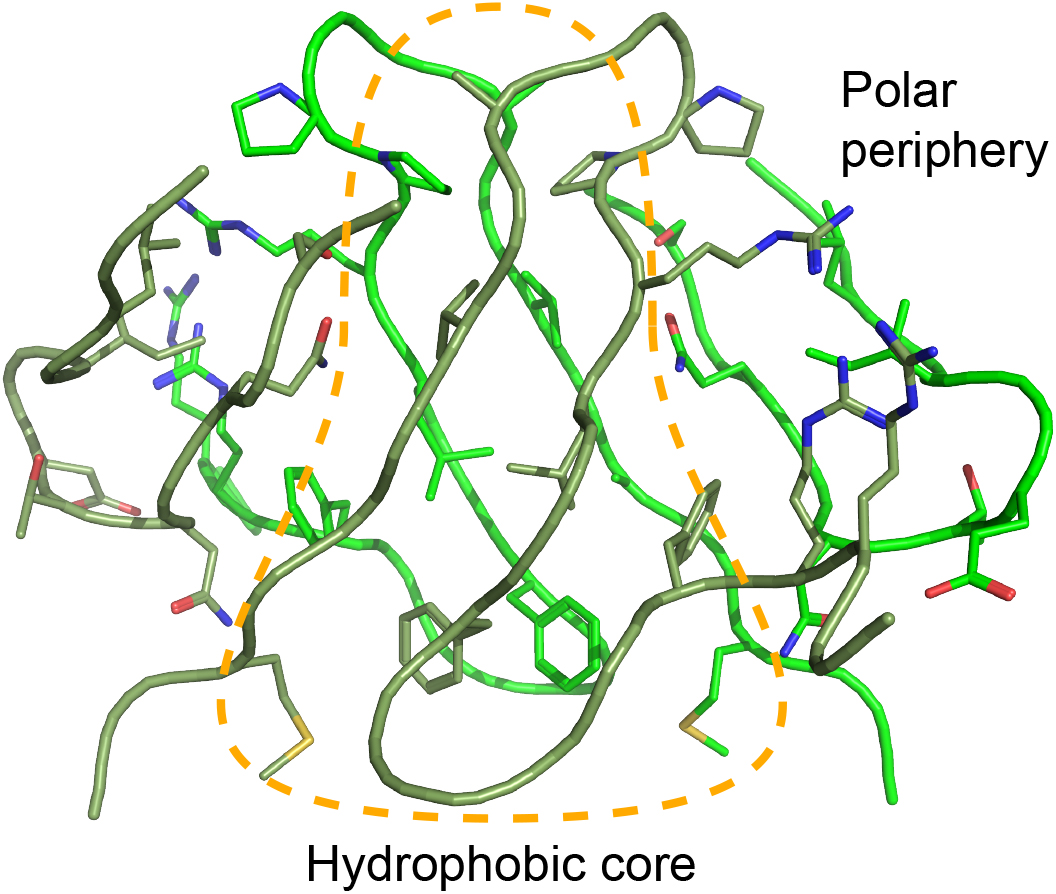
The RIG-5 homodimeric interface has a hydrophobic core and polar periphery, as was observed for the Dpr-DIP and DIP-DIP complexes (ref. 8, 10).

**Fig. S6.**
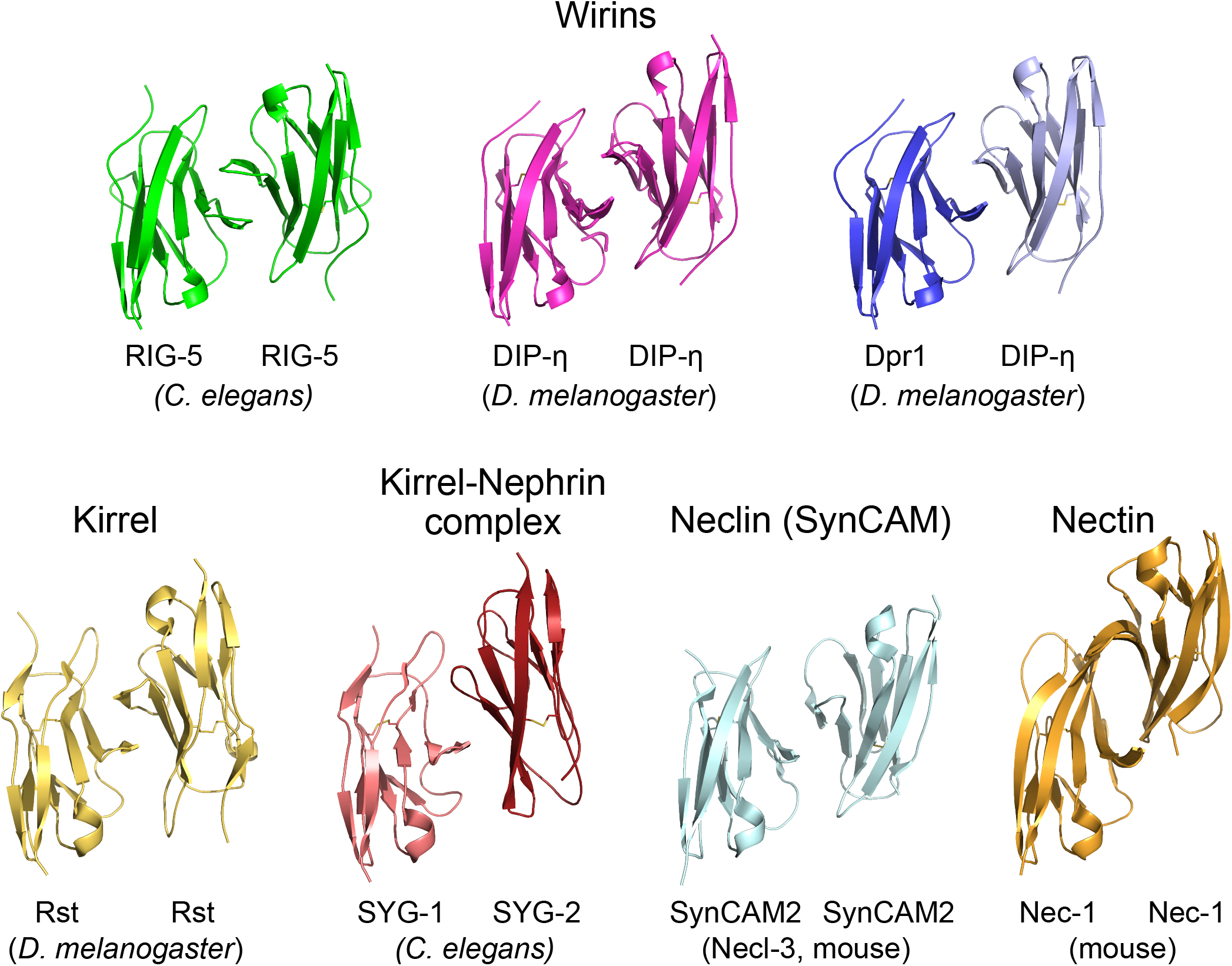
Comparison of IG1-IG1 complexes from Wirins and related families. Family names are noted above the structures. The structures were aligned so that the subunits depicted on the left are superimposed on to each other. The PDB IDs of the structures shown are XXXX, XXXX, XXXX, 4OF8, 4OFY, 3M45, and 5B21.

